# Identifying conserved molecular mechanisms of thermo-acclimation in symbiotic organisms

**DOI:** 10.1101/492702

**Authors:** H. J. Alves Monteiro, C. Brahmi, A. B. Mayfield, J. Vidal-Dupiol, B. Lapeyre, J. Le Luyer

## Abstract

Seawater temperature rise in French Polynesia has repeatedly resulted in symbiosis breakdown between giant clam (*Tridacna maxima*) and dinoflagellates (*Symbiodinium* spp.), particularly in small individuals. Herein, we explored the physiological and gene expression responses of the clam hosts and their photosynthetically active symbionts over a 65-day experiment in which clams were exposed to either normal or environmentally relevant elevated seawater temperatures. These data were combined with publicly available data for both free-living *Symbiodinium* (clades C1 and F) and *Symbiodinium* spp. in *hospite* with the coral *Pocillopora damicornis*. Gene module preservation analysis revealed that the function of the symbionts’ photosystem II was impaired at high temperatures, and this response was conserved across all holobionts and *Symbiodinium* clades examined. Similarly, activation of the phytohormone abscisic acid signaling and epigenetics modulation appeared to be a key response mechanisms for symbionts in *hospite* with giant clams exposed to high temperatures and also distinguish thermo-tolerant from thermo-sensitive *Symbiodinium* C1 phenotypes.

## Introduction

Giant clams are mixotrophic organisms living in obligatory symbiosis with photosynthetic dinoflagellates of the genus *Symbiodinium* (*1*, *2*). *Symbiodinium* spp. associate not only with giant clams, but with a diverse array of marine invertebrates, namely sponges, molluscs and cnidarians; indeed, the *coral-Symbiodinium* symbiosis is the functional basis of all coral reefs (*3*). While in scleractinian corals symbionts are located intracellularly, in giant clams they reside extracellularly inside a tubular system called “Z-tubules,” which are 1) found in the outer epithelium of the mantle and 2) connected to the stomach (*4*). These *in hospite* dinoflagellates are known to provide nutrients to the clam hosts via photosynthesis and may account for a major part of the clams’ energy needs (depending on the species and the life history stage) (*5*–*9*). The *Symbiodinium* genus includes nine different clades with well characterized molecular and physiological differences (*10*–*12*). One *Symbiodinium* clade, clade A (*S. microadriaticum* and *S. fitti*), has been recurrently found in symbiosis with *Tridacna maxima* though members of clades C and D have been found in clam tissues, as well (*10*, *13*–*17*). Depending of the species, cladal representation of symbiont types have been found to vary with individual size (mostly observed in *T. squamosa*), as well as across environmental gradients (especially seawater temperatures) (*15*, *16*). In French Polynesia, eastern Tuamotu’s archipelagos were historically characterized by high densities of small giant clams (*18*–*20*). Recent mortality episodes and/or “bleaching” events in the Tuamotu Islands have, however, been reported, including 1) a massive mortality event in 2009 that reduced the small giant clam population by 90% at Tatakoto Atoll (*18*, *21*) and 2) a bleaching event in 2016 that affected 77 and 90% of the wild and cultured giant clam populations, respectively, at Reao Atoll (*22*). An increase in surface seawater temperature over a prolonged period (approximately three months above 30°C) is suspected to have triggered such bleaching events (*18*, *21*, *22*).

As with corals, bleaching in small giant clams corresponds to the loss of symbiotic *Symbiodinium* from the hosts (*18*, *23*–*26*)*. Symbiodinium* spp. community variability and diversity (i.e., the representation of various clades and/or sub-clades) seems to be a determining factor in the sensitivity and the resilience of both coral and small giant clam hosts to increased temperatures (*27*–*30*). However, the cell physiology of the host and symbionts is likely to be as important, if not more so, than the *Symbiodinium* assemblage, in terms of gauging the ability of the clam-*Symbiodinium* symbiosis to acclimate to elevated temperatures over prolonged durations. To date, only a few studies have investigated the transcriptomic response of giant clams to elevated temperatures while lipid profiling analyses are more routinely undertaken (*31*). The transcriptomic response to elevated temperature of several other taxa, mostly scleractinian coral species (*32*–*35*) and cultured *Symbiodinium* spp. (*36*, *37*) have also been explored, yet few studies have looked at the mRNA level responses of multiple *Symbiodinium* clades and host systems in the same study. Furthermore, few physiological data and even fewer transcriptomic data are available for the high-temperature responses of the giant clam *T. maxima* and its symbionts (but see (*31*, *38*)); these two published studies, though, only considered the response to an abrupt, rapid (and environmentally unrealistic) increase in temperature. Consequently, our understanding of the possible key drivers in high-temperature acclimation remains largely incomplete, despite its importance for generating better predictions of the impact of climate change on wild populations of giant clams (*21*). Given such knowledge deficiencies, we aimed herein to characterize the physiological and transcriptomic responses of small giant clams and their symbionts to sub-lethal elevated temperatures (~30.7°C) over a two-month period. In addition to hypothesizing that the giant clams would ultimately acclimate to this experimentally elevated temperature, we further hypothesized that a “dual-compartmental” bioinformatic approach, similar to the one that has been used with corals (*39*) would provide insight into the key molecular pathways underlying the ability of each member of this association to acclimate to an environmentally relevant, sub-lethal temperature.

## Results

### (a) Physiology

We observed no mortality across the 65-day experiment, but some of the individuals exposed to elevated temperatures showed signs of partial bleaching in the 30.7°C treatment by day 65. *Symbiodinium* density and photosynthetic yield (Fv/Fm) were both lower in clams exposed to elevated temperatures (Scheirer-Ray-Hare; H=24.44, *p<*0.001 and H=22.88, *p<*0.001, respectively; Supplementary figure S1). There was no interaction between time and temperature for either of these response variables, and *Symbiodinium* Fv/Fm remained constant over the three sampling times (Scheirer-Ray-Hare; H=1.26; *p*=0.53, Supplementary figure S1). Time had only a slight effect on *Symbiodinium* spp. density (Scheirer-Ray-Hare; H=6.07; *p*=0.048, Supplementary figure S1), though no *post-hoc* differences were detected between individual sampling times (Dunn’s test; alpha = 0.05).

### (b) *Symbiodinium* spp. communities in clams

The *Symbiodinium* spp. communities of all small giant clam hosts (from both control and high temperature conditions) were primarily composed of clade A (>99%). Four clams showed secondary populations of clade C (ranging from 1.8 to 32.8% of the total cell population), as well as residual quantities of clades B and F (<0.001%). There were no detectable effects of prolonged high-temperature exposure of the *Symbiodinium* assemblages within the giant clam samples (Supplementary figure S2). *In situ* clam samples from Reao atoll (Tuamotu Archipelago, French Polynesia) confirm predominance of clade A (mean 93.0% ± 10.7) with presence of minor clades B + C (mean ~5 %) and suggest that transport and experimental facility environment did not impact relative cladal representation in this study (Supplementary data).

Metabarcoding internal transcribed spacer 2 (ITS2) sequencing resulted in an average of 186.7k ± 25.7 PE sequences per sample. After sequences pre-processing, Dada2 algorithm reported a total of 12 amplicon sequence variants matching to clades A (N=9) and C (N=3) that parallel results from qPCRs (Supplementary figure S2, C). Clade A sequence variants mainly match to sub-clade Clade A3 (best-hit BLASTn; *e*-value<10^−6^). Clade representation based on UniFrac distance (PERMANOVA; pseudo-F=1.3; q-value=0.33) or evenness values (Kruskall-Wallis; H=0.04; q-value=0.83) did not significantly differ between temperature conditions.

### (c) Transcriptome assemblies

A total of 363.70 million 100-bp paired-end reads were used to assemble a raw meta-transcriptome (host + symbionts) of 726,689 transcripts (420.02G bp). After stringent filtering and segregation of host and *Symbiodinium* spp. sequences, the assemblies resulted in a transcriptome for *T. maxima* of 24,234 contigs (N50=1,011 bp; GC content=40.1%) and a meta-transcriptome for *Symbiodinium* spp. of 51,648 contigs (N50=1,027 bp; GC content=57.9%). High G-C content is generally a hallmark of *Symbiodinium* spp. transcriptomes (*40*). Transcriptome statistics and annotations are provided in Table 1 and Supplementary table S1, respectively.

**Table 1:** Transcriptome assembly statistics. (A) Table showing main assembly metrics for *Tridacna maxima* and *Symbiodinium* spp. (B) Density plot of the percent G-C content for *Symbiodinium* spp. and *Tridacna maxima* contigs.

| *Tridacna maxima* |  |
| --- | --- |
| Total number of transcripts | 24,234 |
| Average percent G-C | 40.15 |
| Contig N50 | 1,768 |
| Median contig length (bp) | 1,011 |
| Average contig length (bp) | 1,276.13 |
| Total assembled bases | 30,925,845 |
| *Symbiodinium* spp. |  |
| Total number of transcripts | 51,648 |
| Average percent G-C | 57.9 |
| Contig N50 | 1,027 |
| Median contig length (bp) | 688 |
| Average contig length (bp) | 845.89 |
| Total assembled bases | 43,688,343 |

### (d) Host clam acclimation response to prolonged high-temperature exposure

A gene co-expression network was built using the normalized RNA-seq data from which low-expression genes had been eliminated, and five modules correlated significantly (*p<*0.05) with temperature, clade composition, and/or physiological data (including oxygen production, *Symbiodinium* spp. density, photosynthetic rate, and host dry weight; Figure 1). No module was correlated with sampling time, O_2_ consumption, or shell extension. Two host modules positively (R=0.82) and negatively (R=-0.46) correlated with temperature, namely the pink_hos_t and the magenta_hos_t modules, respectively (Figure 1). The magenta_hos_t module also correlated positively with photosynthetic rate Fv/Fm (R=0.63) while the pinkh_os_t module correlated negatively with it (R=-0.51). Only the pinkh_os_t module also negatively correlated with oxygen production (R=-0.44; Figure 1). Among the most enriched GO terms in the pinkh_os_t module were secondary metabolic processes (GO:0019748), pituitary gland development (GO:0021983), locomotion (GO:0040011), and cilium movement (GO:0003341; Supplementary table S2). The magenta_hos_t module was enriched for carbohydrate biosynthetic processes (GO:0016051), biological adhesion (GO:0022610), phenol-containing compound metabolic processes (GO:0018958), cofactor metabolic processes (GO:0051186), and response to stimuli (GO:0050896; Supplementary table S2).

**Fig. 1:**
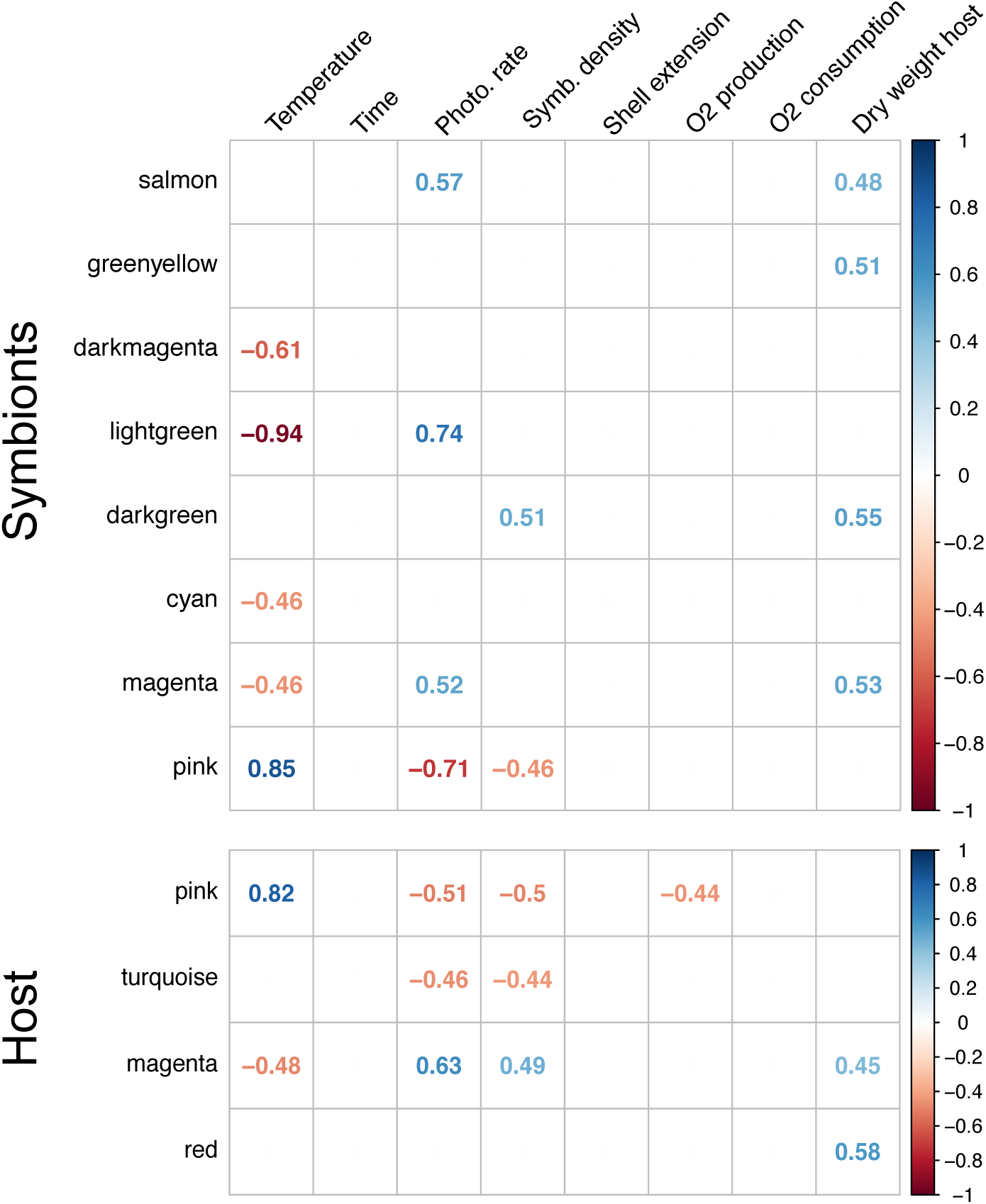
Correlation matrix of symbionts and host gene expression against condition factors and quantitative physiological traits. Genes have been clustered in modules (y-axis) according to their co-expression values. Values in cells indicate Pearson’s correlation scores, and only statistically significant correlations (*p<*0.05) are depicted.

### (e) *Symbiodinium* acclimation to prolonged high-temperature exposure

Co-expression network analysis of *Symbiodinium* spp. showed more modules correlated with temperature than for the clam host, either negatively (darkmagenta_symbiont_ [R=-0.61], lightgreen_symbiont_ [R=-0.94], cyan_symbiont_ [R=-0.46], and magenta_symbiont_ [R=-0.46]) or positively (pink_symbiont_ [R=0.85]). Among the enriched GO terms in the pink_symbiont_ module were RNA metabolic processes (GO: 0016070), methylation (GO:0032259), mitochondria-nucleus signaling pathways (GO:0031930), and response to abscisic acid (GO:0009737). For the magenta_symbiont_ module, enriched genes tended to be involved in glucan biosynthetic processes (GO:0009250), movement of cells or sub-cellular components (GO:0006928), energy derivation by oxidation of organic compounds (GO:0015980), and sodium ion transport (GO:0006814). The darkmagenta_symbiont_ module was enriched for photosynthetic electron transport chain (GO:0009767), oxidation-reduction processes (GO:0055114), regulation of the response to stimuli (GO:0048583), and chloroplast thylakoid membrane (GO: GO:0009535). Complete GO enrichment results can be found in Supplementary table S2.

We further identified the top 30 most 1) “connected” (i.e., high connectivity determined by the co-expression network analysis) and 2) annotated genes (hereafter referred as “hub genes”) within each relevant module. For the pink_symbiont_ module, the hub genes were a putative D-lactate dehydrogenase, mitochondrial LDHD, photosystem II reaction center protein L (PSBL), Ras-related protein Rab-11A (RAB11A), phosphoenolpyruvate/phosphate translocator 3, chloroplastic PPT3, and the cytochrome c biogenesis ATP-binding export protein CcmA (CCMA). For the darkmagenta_symbiont_ module, we identified a choline transporter-like protein 5 (SLC44A5), D-alanine aminotransferase (DAT), cAMP-dependent protein kinase type I-alpha regulatory subunit (PRKAR1a), kinesin-like protein KIF3A (KIF3A), non-specific lipid-transfer protein 1, photosystem I P700 chlorophyll a apoprotein A1 (PsaA), and the NFX1-type zinc finger-containing protein 1 (ZNFX1). Exhaustive lists of hub genes for each module are provided in Supplementary table S3.

We tested for specific module conservation across datasets for the *Symbiodinium* spp. transcriptomic response in the small giant clams. Overall, few modules were conserved between datasets (Figure 2). We observed that the pink_symbiont_ module was moderately conserved in free-living *Symbiodinium* spp. (Zsummary=2.9) but not in the coral datasets (Zsummary=-0.5). Whereas the magenta_symbiont_ tended to be conserved in corals (Zsummary=1.9), it was not conserved in free-living individuals (Zsummary=1.3). Interestingly, module lightgreen_symbiont_ was not conserved across datasets despite being the *Symbiodinium* spp. module that was most strongly correlated with temperature in small giant clams (Figure 1). One module only, the darkmagenta_symbiont_, was, however, preserved across all datasets (Zsummary=4.1 and 7.6 for *coral-Symbiodinium* and cultured *Symbiodinium* datasets, respectively; Figure 2). As mentioned above, this module was enriched for genes involved in photosynthetic activity and maintenance.

**Fig. 2:**
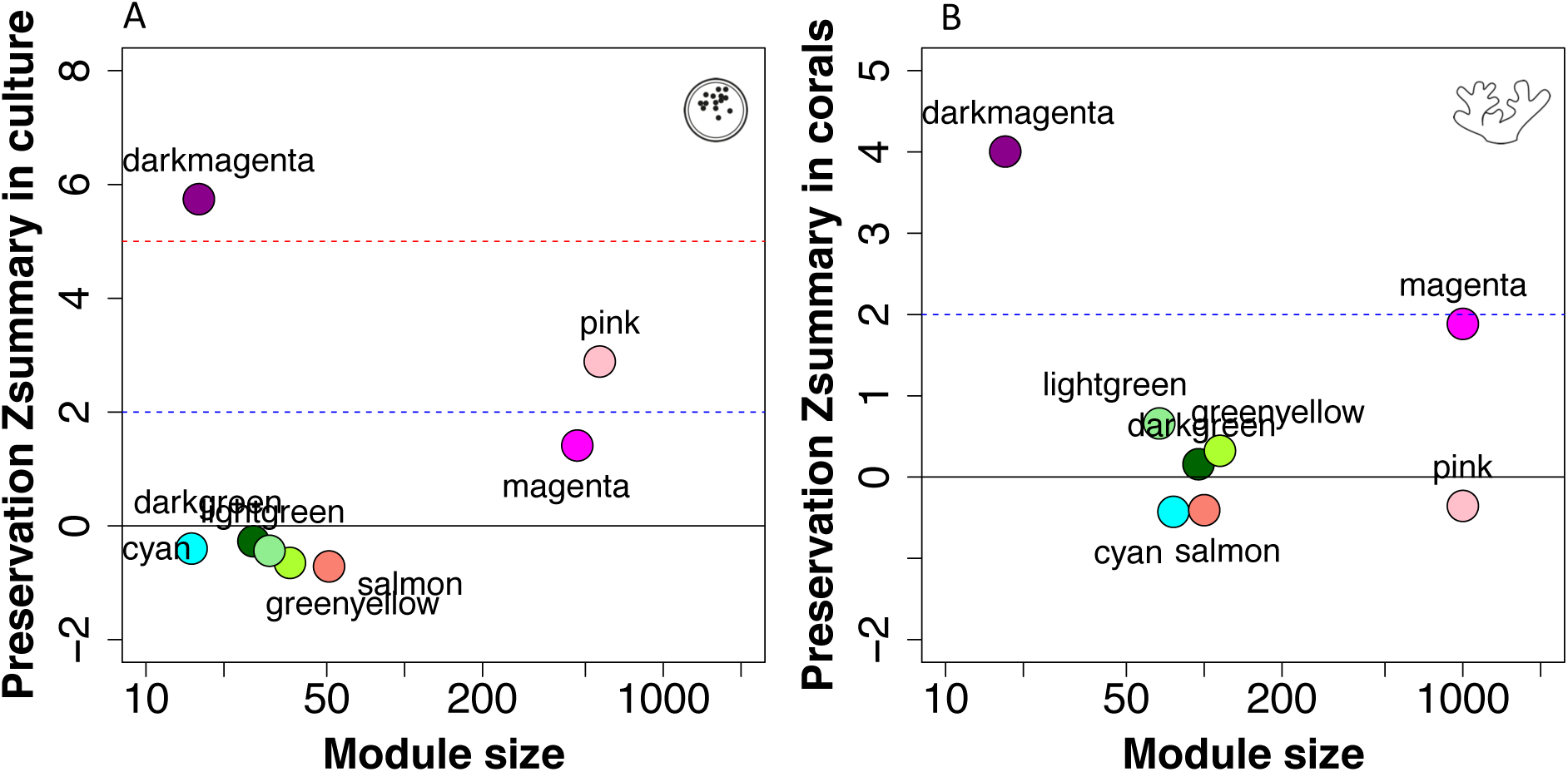
Module preservation statistics. (A) Zsummary values plotted against module size (i.e., number of genes in the module) for the conservation of modules identified in symbionts within clam hosts against free-living *Symbiodinium* spp. (B) Zsummary values plotted against module size for the conservation of modules identified in symbionts within clam hosts against *Symbiodinium* spp. symbionts within the coral host *Pocillopora damicornis*.

### (f) Multivariate analysis and co-expression network for free-living datasets against the clade C1 reference genome

We used db-RDA to document gene expression variation among free-living *Symbiodinium* spp., with temperature and cladal breakdown as the explanatory variables. The model was highly significant (*p<*0.001), and the adjusted R^2^ was 0.80 (Figure 3). Partial db-RDAs showed that the transcriptomic response was largely lineage-specific (adj. R^2^=0.77; F=179.88, *p*<0.001). However, temperature also had a significant effect on total gene expression variation, independent of *Symbiodinium* genotype (adj. R^2^=0.05; F=12.42, *p*<0.001). We conducted independent WGCNA analyses to assess acclimatory responses in free-living *Symbiodinium* spp. based on genome C1 specific (*41*). We found the largest modules (blue (n= 8,739 genes) and turquoise (n = 10,061 genes) to be correlated with lineage (Supplementary figure S3), independent of the temperature response; this is congruent with the db-RDA analysis (see Supplementary material for details.). Similarly, we found two modules (brown_C1_ and red_C1_) that were significantly correlated with temperature (R=0.82 and -0.89, respectively; Supplementary figure S3). Finally, we found a single module (green_C1_; n = 123 genes) positively correlated with thermotolerance (R=-1); it could effectively segregate thermo-sensitive *Symbiodinium* from thermo-tolerant phenotypes. Among the most enriched GO terms we found DNA methylation (GO:0006306), genetic imprinting (GO:0071514), suggesting profound methylation changes between phenotypes. Furthermore, several ancestral GO terms linked to abscisic acid-activated signaling including response to acid chemical (GO:0032776), cellular response to alcohol (GO:0097306), response to endogenous stimulus (GO:0009719) and cellular response to oxygen-containing compound (GO:1901701) were among the most enriched functions (Supplementary figure S3 and table S2).

**Fig. 3:**
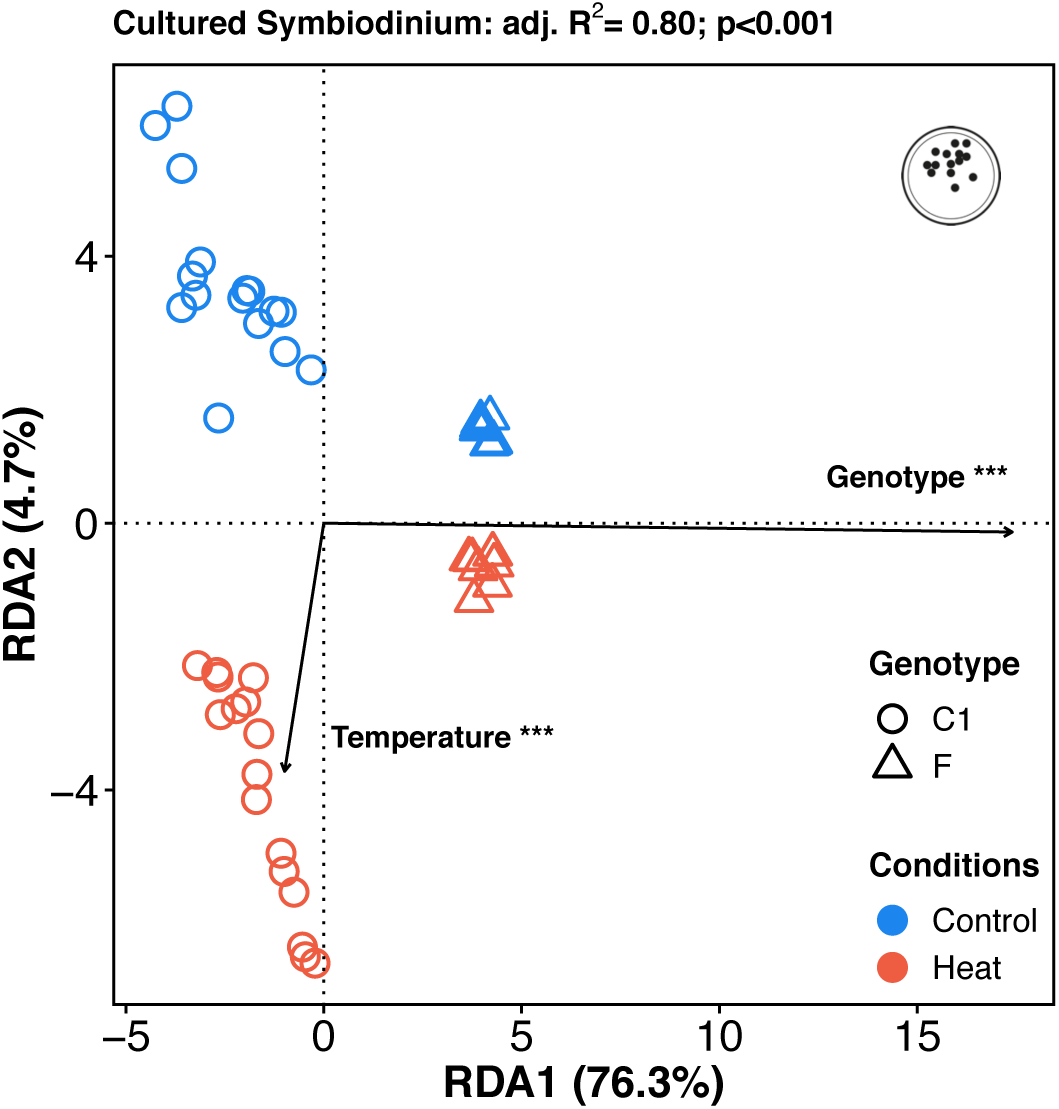
RDAs of cultured *Symbiodinium* (clades C1 and F). The overall model was statistically significant (*p<*0.001) and accounted for 80.0% of the variance explained. “***” indicates *p<*0.001, (effect of explanatory factor).

## Discussion

Temperature increases are threatening marine invertebrate populations worldwide, especially for species already living at, or close to, their upper thermal tolerance limits (*42*). Recent heat wave events have resulted in ~90% declines in *T. maxima* populations in some atolls of French Polynesia (*18*, *22*). While several studies have investigated the invertebrate (molluscs and cnidarian) response to heat stress over short-term timescales, relatively few have investigated the prolonged response to elevated temperatures (*39*). Although our small giant clam samples ultimately acclimated to an experimentally elevated temperatures of nearly 31°C, *Symbiodinium* density was reduced in thermally challenged clams, and both host clams and their *Symbiodinium* spp. populations underwent gene expression changes over the course of this two-month experiment. Upon discussing such temperature-driven changes in gene expression, we highlight some intrinsic responses of the symbionts (i.e., independent of the host species) and identify key mechanisms involved in thermal-tolerance in the symbionts.

### (a) Symbiosis specificity might preclude symbiont shift as a thermal acclimation strategy in *T. maxima*

A 1.5°C temperature elevation over a 65-day period was sufficient to induce a significant reduction in symbiont density in small giant clams; no bleaching (even partial) was observed in control temperature clams. Our results support previous studies of corals and giant clams in which high temperature exposure led to sub-lethal bleaching (*25*, *38*, *43*–*47*); whether the cellular mechanisms of bleaching are conserved between corals and giant clams remains to be determined (*38*, *48*). For some coral species, resilience to heat stress is associated with a more flexible symbiotic association (i.e., the capacity to shift from one dominant *Symbiodinium* clade to another) (*49*–*53*). Indeed, some bleaching events have largely been attributed to the thermal sensitivity of specific endosymbiotic *Symbiodinium* spp. residing in coral host tissues (*54*, *55*). Corals hosting *Symbiodinium* clade C is more prone to bleaching, whereas those housing certain lineages of clade D *Symbiodinium* have demonstrated an improved thermotolerance (*56*, *57*). Similarly, clade C and/or D are more commonly found in giant clams inhabiting warmer environments while clade A *Symbiodinium* are more common in clams located in cooler waters (*15*). Herein, the *Symbiodinium* spp. communities were predominantly clade A, even after two months of high temperature exposure; this finding aligns with other studies in corals that found *Symbiodinium* assemblages to be temporally stable, even as environmental conditions changed (*58*–*61*). This was not an artifact due to our experimental conditions since individuals sampled from their original location also host predominantly clade A.

Such a high proportion of clade A in giant clams is expected and it has also been reported in the sea anemone *Anemonia viridis*, while it is in sharp contrast to other hosts such as corals, which host a broader clade diversity (*62*–*64*). This near-exclusive hosting of clade A in clams, and the temporal stability of their association, suggests that some selection process favor clade A (or else impairs recruitment of other clades); lectin/glycan interactions surely playing a role in such recognition-related processes (*65*). Admittedly, broader *in situ* data encompassing different time of the year, will be necessary to verify the fidelity between clade A *Symbiodinium* and giant clams, and whether mixed-clade assemblages are common *in situ* (*66*). Nevertheless, such reduced flexibility would preclude community shift as a strategy for the small giant clam to cope with increased temperatures, at least in our experimental context. Rather than adaptation (i.e., a community shift resulting in a new “holobiont genomic landscape”), acclimation (i.e., physiological changes that initially manifested at the molecular level) appears to have played a larger role in this study.

### (b) Effect of prolonged exposure to elevated temperature on the small giant clam transcriptome

Both host and *Symbiodinium* spp. gene expression were affected by elevated temperature exposure, with no significant effects of time from 29 days onwards; the temperature-related differences were from thenceforth sustained over time. We found one gene module positively impacted by temperature and negatively correlated with symbiont photosynthetic yield (Fv/Fm) and density. This module was enriched for tryptophan metabolic processes and included the tryptophan 2,3-dioxygenase coding gene (TDO), a pivotal regulator of systemic tryptophan levels also involved in the response to oxidative stress (*67*, *68*). Tryptophan is the precursor of 5-hydroxytryptamine (5-HT), a bivalve serotonin transmitter that plays critical roles in numerous physiological functions (e.g., reproduction (*69*)). In the coral *Montastrea faveolata* (now known as *Orbicella faveolata*), TDO (called AGAP in the reference [94]) was up-regulated in response to ultraviolet radiation in larvae and was associated with individual fitness (locomotion and settlement) impairment in later stages (*70*).

The negatively impacted module was enriched for glyceraldehyde-3-phosphate metabolic processes and was correlated with higher photosynthetic yield (Fv/Fm). This gene, as well as a large number of others encoding proteins involved in metabolic processes, were also significantly affected by a sub-lethal elevated temperature (30°C) in the reef coral *P. damicornis* (*39*). Pollutant exposure also altered the expression of genes involved in carbohydrate metabolism, albeit only in the coral host compartment (and not in *Symbiodinium*) in another study (*71*). Admittedly, we did not assess the proportion of the energy derived from autotrophy, which ranges widely (from 25 to up to 100%) and is dependent on the species and/or life history stage in the *Tridacna* genus (*6*, *7*, *72*); shifts from autotrophy to heterotrophy, and vice versa, are likely to affect host gene expression patterns. However, the facts that 1) carbohydrate metabolism plays a central role in maintenance and stability of the clam-dinoflagellate holobiont (*73*) and 2) this gene module was correlated with photosynthetic yield and symbiont density provide support for the existence of a mechanism by which the host influences symbiont performance. All together, our results suggest that the combined effects of regulation of tryptophan levels and impairment of carbohydrate metabolism might be key elements in the long-term response to elevated temperature in small giant clams. However, how these changes would affect fine-scale interactions between the host and symbionts remains to be explored; hence further proteomic data that directly speak to changes in cell physiology should be acquired in future studies of giant clam*-Symbiodinium* symbioses.

### (c) *Symbiodinium* spp. response to prolonged elevated temperature exposure *in hospite*

Overall, *Symbiodinium* spp. gene expression decreased with prolonged elevated temperature exposure, and some of the modules were correlated with the lower *Symbiodinium* photosynthetic yield and cell densities documented at elevated temperatures. Physiological studies have previously shown that high temperature led to diminished photosynthetic yield in several clades of *Symbiodinium* spp. (*74*). Herein we also found that genes encoding certain components of the photosynthetic machinery, especially photosystem II (PSII), were tune down in elevated temperature condition. PSII integrity is vital for proper *Symbiodinium* function, and PSII damage has been directly linked to bleaching in corals (*45*). It is noteworthy that the same gene module also included chloroplast thylakoid membrane rearrangement-related genes, which are used by *Symbiodinium* spp. and other photosynthetic organisms to cope with heat and high UV radiation (*75*, *76*). Although the small giant clams generally appear to have acclimated to elevated temperatures over our two-month experiment, the *Symbiodinium* communities may, instead, have been manifesting signs of intracellular stress given these gene expression changes, as well as the decreases in cell density and Fv/Fm. Whether or not these holobionts could have sustained an even longer exposure to 31°C remains to be determined, though it is worth noting that, unlike *in situ*, clams were not fed in the aquaria. It is thus most likely that clams allowed to feed both autotrophically and heterotrophically might, then, have an even superior capacity for high-temperature acclimation.

Although the effects of clade and/or experiment on *Symbiodinium* gene expression were much greater than the temperature effect, temperature significantly impacted gene expression even when controlling for clade genotypes. This suggests that acclimation to a long-term temperature increase has a conserved basis among different lineages despite large evolutionary distance between clades (*10*, *77*). Differences in gene expression between clades might also be associated with distinctive thermotolerance capacities. We found that few gene modules were preserved across datasets: *in hospite* with clams vs. free-living vs. *in hospite* with corals. One methodological factor that may warrant a cautious interpretation of our results is the major experimental design differences between datasets; in the case of our own study, as well as that of Mayfield *et al*. (*39*), sub-lethal temperatures were used, whereas in exposure time were more reduce in Levin *et al*. (*37*) and Gierz *et al*. (*36*) and it is most likely that individuals were still experiencing stress. However, the fact that we were still able to detect a module related to photosynthesis in all studies suggests that certain molecular pathways underlying the photosynthetic activity of *Symbiodinium* are temperature sensitive, regardless of whether the *Symbiodinium* are free-living or associated with corals or giant clams.

### (d) Evidence for thermotolerance mechanisms in *Symbiodinium* spp

Exposure to elevated temperature was associated with an up-regulation of phytohormones and methylation-related genes. A single gene module associated with profound methylation changes revealed higher gene expression with elevated temperature and was negatively correlated with *Symbiodinium* density and photosynthetic yield. It is known in plants that DNA methylation and histone modification are associated with the response to heat stress, and, more specifically, act to prevent heat-associated macromolecular damage (*78*). Such methylation changes might be inherited and account for, in part, the remarkable ability of plants to adapt and/or acclimate quickly to stressful environments (*79*, *80*). Additionally, studies in other plant systems found that phytohormone ABA and reactive oxygen species (ROS) regulation are key molecular mechanisms involved in the capacity to acclimate to abiotic stressors including oxidative stress tolerance in unicellular algae (*81*). Indeed, an increase in ABA biosynthesis is associated with up-regulation of ABA signaling genes in several plant species (*82*). Herein we showed that the same gene module enriched for methylation regulation was also enriched for ABA signaling/metabolism, and this response was conserved in free-living *Symbiodinium* cells. Additionally, independent data from thermo-sensitive and thermo-tolerant phenotypes of free-living *Symbiodinium* (clade C1) revealed that the only biological function associated with thermal tolerance was the regulation of methylation and response to acid chemical most likely related to ABA signaling. Considering the pivotal role of ABA in a variety of plant systems, we propose that ABA signaling might also be a key driver of acclimation to long-term elevated temperature exposure, and, more generally, thermal tolerance in *Symbiodinium* spp. Ultimately, tight co-expression between ABA-responsive genes and methylation effectors also suggests a possible epigenetic control of ABA signaling during thermal acclimation in *Symbiodinium* spp. that remains to be elucidated.

## Conclusion

The co-expression network analysis proved to be a powerful tool for dissecting compartment-specific transcriptomic responses in symbiotic systems. This is especially true when looking for acclimatory signatures that, in contrast to short-term stress responses, are characterized by rather subtle changes over longer periods. Indeed, our data from a long-term high temperature study revealed that different cellular processes are impacted in the host clam and *in hospite Symbiodinium* compartments; genes encoding key photosynthesis proteins were particularly temperature sensitive in not only *Symbiodinium in hospite*, but also in culture. Future studies focusing on the range of optimal thermal conditions of the *T. maxima* species may improve our understanding on the thermal tolerance of the small giant clams and their symbionts. Although the giant clams used in this study ultimately survived a two-month exposure to nearly 31°C, it is possible that slightly higher temperatures or extended exposure might cause them to bleach to such a great extent that they would not survive. Herein, we lifted the veil on a novel mechanism involving epigenetic landscape rearrangement and phytohormone regulation that provide *Symbiodinium* with improved thermotolerance capacities. How the impact of stressful environmental conditions might be impacting the subsequent generation’s tolerance and/or physiological capacities (i.e., epigenetic effects) must be addressed in the near future.

## Materials and Methods

### (a) Experimental design, tissue sampling, and physiological measurements

Experimental procedures were first described by Brahmi et *al*. (*43*). Briefly, a total of 24 individual small giant clams (N=4/treatment) were sampled over a 65-day period (days 29, 53, and 65) in control (29.2°C; ambient at the time of experimentation) and elevated (30.7°C) temperature conditions (N=4 tanks/treatment). The temperatures employed and the duration of the experiment reflect conditions in normal and abnormally hot seasons reported in French Polynesia Tuamotu’s lagoons (*43*), respectively, and field bleaching of Tuamotu small giant clams has been previously documented when temperatures are sustained at >30°C for five months with peak at 31.8°C (*83*). Yet, optimal temperature and critical point values remain to be precisely assessed for giant clams.

Samples (approx. 1 cm^2^) from each of the two treatments at each of the three sampling times were systematically collected from the same region of the mantle and stored in RNA-Later^®^ (Life Technologies, USA) at -80°C until analysis (N=24). Furthermore, a single hermaphroditic individual (approximately two years old) was sampled for a total of seven different tissues (mantle, adductor muscle, gonads, gills, byssus, visceral mass, and kidney) for transcriptome assembly. Only one individual was used in an effort to reduce assembly polymorphism biases. For this individual, sexual status was confirmed by gonad biopsy and histology following a previously detailed procedure (*84*). This individual was not included in the differential expression analysis. Additionally, 10 giant clams were collected *in situ* in October 2018 in the Reao atoll (Tuamotu Archipelago, French Polynesia). Samples (approx. 1 cm^2^) from each of the samples was collected from the same region of the mantle and stored in 95% ethanol at -20°C until later symbionts community analysis.

As described in detail in Brahmi et *al*. (*43*), a variety of physiological response variables were assessed in the 24 experimental replicates, in addition to the profiling of their transcriptomes: growth, *Symbiodinium* density, and the maximum dark-adapted yield of photosystem II (Fv/Fm; as measured by an AquaPen pulse amplitude modulator fluorometer (APC-100m, Photon System Instruments, Tchek Republic). Please see our prior work for details on these analyses. Physiological data were tested with two-way ANOVA (treatment x time) followed by Tukey’s “honestly significant difference” (HSD) *post-hoc* tests (*p<*0.05), including the interaction between time and temperature when data, or transformed data, met the assumptions for ANOVA. For *Symbiodinium* density and Fv/Fm, a non-parametric equivalent of the two-way ANOVA, the Scheirer-Ray-Hare test, was instead used; with post-hoc Dunn’s test.

### (g) DNA/RNA extractions and transcriptome sequencing

Total RNA was extracted from *T. maxima* mantles by lacerating tissues with a scalpel and rinsing with 1X PBS. Cellular lysis was induced by addition of 1.5 ml TRIzol (Invitrogen, USA) according to the manufacturer’s recommendations. The supernatant was transferred into a 2-ml tube and incubated for 10 min on ice. The phase separation was achieved by addition of 300 μl of chloroform coupled with centrifugation at 12,000 *xg* for 12 min at 4°C. The upper aqueous layer contained the RNA, and the lower organic layer was stored at -20°C for later DNA extraction (according to the manufacturer’s recommendations). Total RNA from each individual was subjected to a DNAse treatment using Qiagen’s RNA cleanup kit (Germany). RNA and DNA were quantified using a NanoDrop ND-2000 spectrophotometer (Thermo-Fisher, USA), and RNA quality was further evaluated by a Bioanalyzer 2100 (Agilent, USA). High-quality RNA was sent to McGill University’s “Genome Quebec Innovation Center” (Montréal, QC, Canada) for Nextera XT (Illumina; USA) library preparation and sequencing on an Illumina HiSeq4000 100-bp paired-end platform. Samples for transcriptome assembly (N=7) were sequenced on a single lane, while samples for differential expression analysis (N=24) were uniformly and randomly distributed over two sequencing lanes after barcoding.

### (h) Transcriptomes assembly

Raw reads provided by RNA-Seq were filtered for quality and length using Trimmomatic v.0.36 (*85*) with minimum length, trailing, and leading quality parameters set to 60 bp, 20, and 20, respectively. Illumina’s adaptors and residual cloning vectors were removed via the UNIVEC database (https://www.ncbi.nlm.nih.gov/tools/vecscreen/univec/). Paired-end filtered reads were assembled *de novo* using Trinity v2.6.6 (*86*) with a default k-mer size of 25 bp and minimum transcript length of 200 bp. Raw transcripts (n=726,689; 420 Gbp) were filtered for presence of open reading frames (ORFs) (length≥300 bp), longest isoform matches, and mapping rate (≥0.5 transcripts per million; TPM).

Transcripts matching Refseq entries from archaea, plasmid, virus, and bacteria (BLASTn; e-value<10^−10^), as well those transcripts that aligned significantly (*e*-value<10^−4^) only to bacterial sequences on the NCBI nt database (max target seqs=5) were discarded in an effort to reduce putative contamination. To segregate between symbiont and host sources, the meta-transcriptome was blasted (BLASTn; *e*-value<10^−4^) against a pool of *Symbiodinium* spp. genomes and transcriptomes including clades A, C, and F (*40*). By default, all hits with no match were considered as originating from the host. For quality checks, the *de novo* transcriptome’s completeness was assessed with BUSCO’s v2 metazoa and v2 eukaryotes databases for clam and *Symbiodinium* spp., respectively (*87*). Transcriptomes were annotated by BLAST search against the Uniprot-Swissprot database (BLASTx; *e*-value<10^−4^). A schematic representation of the overall analysis pipeline has been provided in the Github repository (https://github.com/jleluyer/acclimabest).

### (i) Co-expression network analysis for host and symbionts and functional enrichment of gene modules

Signed co-expression networks were built for the host and symbiont datasets independently using the R package WGCNA following the protocol developed by Langfelder and Horvath (*88*) based on normalized log-transformed expressions values (log2 count per million; log2CPM). The datasets were previously filtered for residually expressed transcripts (>1 CPM in 4 individuals) and minimum overall variance (>10%). Normalization was conducted on library sizes for host and symbiont datasets separately. In an effort to normalize for *Symbiodinium* cells we also tested specific housekeeping genes validated from previous thermal stress experiments (*89*). Since both approaches gave similar results (data not shown), only library size normalization has been presented herein. The main goal of this analysis was to cluster genes in modules correlated with time, temperature, and relevant physiological responses (Figure 1). Briefly, we fixed a “soft” threshold power of 6 and 26 for the host and symbiont datasets, respectively, using the scale-free topology criterion to reach a model fit (|R|) of 0.81 and 0.90 for host and symbionts, respectively. The modules were defined using the “*cutreeDynamic*” function (minimum of 30 genes by module and default cutting-height=0.99) based on the topological overlap matrix, and a module Eigengene distance threshold of 0.25 was used to merge highly similar modules. For each module, we defined the module membership (kME=correlation between module Eigengene value and gene expression values). Only modules with *p<*0.05 were conserved for downstream functional analysis (Figure 1). Gene ontology (GO) enrichment analyses were conducted for each module separately using GOAtools (*90*) implemented with the “go_enrichment” pipeline (https://github.com/enormandeau/go_enrichment) and based on go-basic.obo data (downloaded on May 30, 2018). GO terms were considered enriched at *p<*0.05 (minimum of three genes). GO redundancy reduction was conducted using the “revigo” online tool (*91*).

### (j) Meta-analysis on free-living *Symbiodinium* and coral transcriptomes

We integrated publicly available datasets featuring similar experimental designs (i.e. increased temperature over a long-term timescale) to further unravel the host, symbiont, and holobiont compartments’ responses with respect to high-temperature acclimation. Searches were conducted with the Web of Science platform with this search formula: «symbio* AND RNAseq* AND temperature» together with informal searches with other research engines. A total of three studies met our criteria: Levin et al. (*37*) and Gierz et al. (*36*) for free-living *Symbiodinium* spp. (n=48 transcriptomes) and Mayfield et al. (*39*) for the response of *Symbiodinium* spp. *in hospite* with the scleractinian coral *P. damicornis* (n= 12 transcriptomes). Gierz et al. exposed cultured *Symbiodinium* (clade F) to a 31°C heat stress (control temperature=24.5°C) over a 28-day period (*36*), while Levin et al. exposed clade C1 *Symbiodinium* (both thermo-tolerant and thermo-sensitive sub-types) to a 32°C heat stress (control temperature=27°C) over a 13-day period (*37*). Finally, Mayfield et al. (*39*) exposed corals housing *Symbiodinium* (clade C) to 30°C over a 9-month period (control temperature=27°C), and both the coral hosts and *in hospite Symbiodinium* appeared to have acclimated to this temperature.

Raw data processing and mapping to the newly assembled *Symbiodinium* transcriptome followed the same procedure as described above albeit adapted for single-end reads. In parallel, analysis of free-living datasets was also conducted against the recently published and assembled *Symbiodinium* clade C1 reference genome for comparisons (*41*). Mapping and downstream co-expression network analysis results are detailed in the Supplementary Material and Supplementary figure S3. We assessed module preservation using the *“modulePreservation”* function (200 permutations) implemented in the WGCNA R package based on the Zsummary composite statistic, taking into account preservation of both connectivity and density in a module (*88*). Modules were considered low to moderately conserved for Zsummary values between 2 and 5 and conserved for Zsummary values > 5 (Figure 2).

In order to determine whether genotype (clade), time, and/or temperature had a significant effect on *Symbiodinium* spp. gene expression variation in free-living individuals and to assess the proportion of variance explained by each factor, we used a redundant-discriminant analysis (RDA; Figure 3). Briefly, a principal components analysis (PCA) was first carried out with a Euclidean distance matrix of the log2-transformed expression levels of the 20,449 genes (log2CPM) using the Ape R package *“daisy”* and “*pcoa”* functions (*92*). Because no axis could be selected according to the broken-stick distribution, we selected all axes explaining at least 2.75% of the variation (4 axes explained 69.5% of the total variance.) according to a previously described method (*93*, *94*). A distance-based redundancy analysis (db-RDA) was then undertaken with the gene expression variation explained by these PCA factors (response matrix), with genotype, temperature, and time as explanatory variables. We only retained explanatory factors that accounted for a significant portion of the variability using the “*ordistep*” function implemented in the vegan R package (*95*). Two partial db-RDAs and analysis of variance (ANOVA; 1000 permutations) were conducted to 1) validate whether each factor separately had a significant impact on the variance (in controlling for the second factor) and 2) estimate the variability explained by each factor (“*RsquareAdf*” function). The effect of a given factor was considered significant when *p<*0.05.

### (k) Quantitative PCR- and meta-barcoding-based *Symbiodinium* clade analysis

We evaluated the relative levels of various *Symbiodinium* clades in our giant clam samples using a series of quantitative PCR (qPCR) assays. Amplifications were carried out on AriaMx real-time PCR System (Agilent, USA) using six primer sets optimized for the amplification of nuclear ribosomal 28S in *Symbiodinium* of clades A-F (*96*) following the protocol of Rouzé *et al*. (*62*). The PCRs (25 μL) comprised 12.5 μL of 2X SYBR^®^ Green master mix (Agilent, USA), 10 μL of DNA (previously diluted to 1 ng μL^−1^), and 1.25 μL of each primer (forward and reverse; each at a stock concentration of 4 μM). PCR thermocycling steps include: 1 cycle of pre-incubation of 10 min at 95°C; 40 cycles of amplification: 30s at 95°C, 1 min at 64°C and 1 min at 72°C; and a final step, for melting temperature curve analysis, of 1 min at 95°C, 30s at 60°C and 30s at 95°C. All measurements were made in duplicate, and all analyses were based on the threshold cycle (Ct) values of the PCR products. Ct values were averaged across duplicate samples when the variation was not exceeding 1; otherwise, samples were re-run until delta Ct<1. Similarity in relative clade abundance was assessed using PCA analysis of a Bray-Curtis similarity matrix using Hellinger-transformed data. Db-RDAs and analysis of variance (ANOVA; 1000 permutations) were conducted to identify whether either temperature or time had a significant impact on *Symbiodinium* assemblage, and an alpha level of 0.05 was set *a priori*. To complement data from the experimental individuals, qPCRs were carried out on mantle fragments from 10 wild individuals and seawater samples collected in Reao Atoll (Tuamotu, corresponding to the geographical origin of the experimental individuals; see Brahmi *et al*. (*43*) for details.) in October 2018. Preparation and analyses were performed as described above and in Rouzé *et al*. (*62*).

As a more detailed means of assessing *Symbiodinium* diversity in the 24 small giant clam samples, a meta-barcoding analysis was undertaken following the protocol of (*97*). Briefly, the ITS2 gene was PCR amplified using previously described primers set (*97*) and sequenced at McGill University’s “Genome Quebec Innovation Center” (Montréal, QC, Canada) on a Illumina Miseq 250 bp paired-end platform. The Dada2 algorithm (*98*), implemented in QIIME2 software (*99*), was used to infer sample exact sequences from amplicon data. The reference database was directly imported from NCBI nt repository and trained on the basis of the ITS2 primers following Cunning *et al*. (*97*) recommendations. Detailed protocol, including scripts, has been made available in a Github public repository (https://github.com/jleluyer/acclimabest).

## Acknowledgments

### General

We are grateful to Michel Pahuatini for providing the small giant clams and seawater samples. We thank Claude Soyez and Leila Chapron for their help in animals rearing. We thank Dr. Laetitia Hédouin, for valuable comments on prior versions of the manuscript. We also would like to acknowledge Corinne Belliard for help in DNA preparation and Dr. Yu-Bin Wang for making the *P. damicornis-Symbiodinium* transcriptomic resources openly accessible, as well as designing the afore-cited *P. damicornis-Symbiodinium* transcriptome server. Finally, we thank O. Bichet for her help with the figures.

### Funding

This experiment was made possible by a grant from Labex CORAIL (France), as well as an IFREMER grant (France; Master’s internship to HAM).

### Author contributions

CB and JVD conceived the GECO project, under which tissue samples featured herein were collected. CB and JLL conceived the transcriptomic study. HAM and JLL carried out the laboratory benchwork. HAM, JLL, and CB analyzed the data. HAM, CB, ABM, and JLL wrote the manuscript. All co-authors contributed substantially to revised drafts of the manuscript.

### Competing interests

We declare that we have no competing interests.

### Data and materials availability

Raw sequencing RNA-Seq data for small giant clams featured herein have been made publicly available on the NCBI database (pending creation), and all scripts discussed in the article can be found on Github (https://github.com/jleluyer/acclimabest). Raw meta-barcoding data are available here: (pending creation). Data for free-living *Symbiodinium* spp. have been previously deposited on the NCBI database: Levin *et al*. (*37*)-BioProject NCBI: PRJNA295075, Gierz *et al*. (*36*)-BioProject NCBI: PRJNA342240. Data for *Symbiodinium* spp. from the reef coral *P. damicornis* (*39*) can be found on the NCBI database (Sequence Read Archive: SRR1030692 and BioProject: PRJNA227785), as well as on this modular, interactive website: http://symbiont.iis.sinica.edu.tw/coral_pdltte/static/html/index.html#home.

## Supplementary Materials

**Fig. S1.**
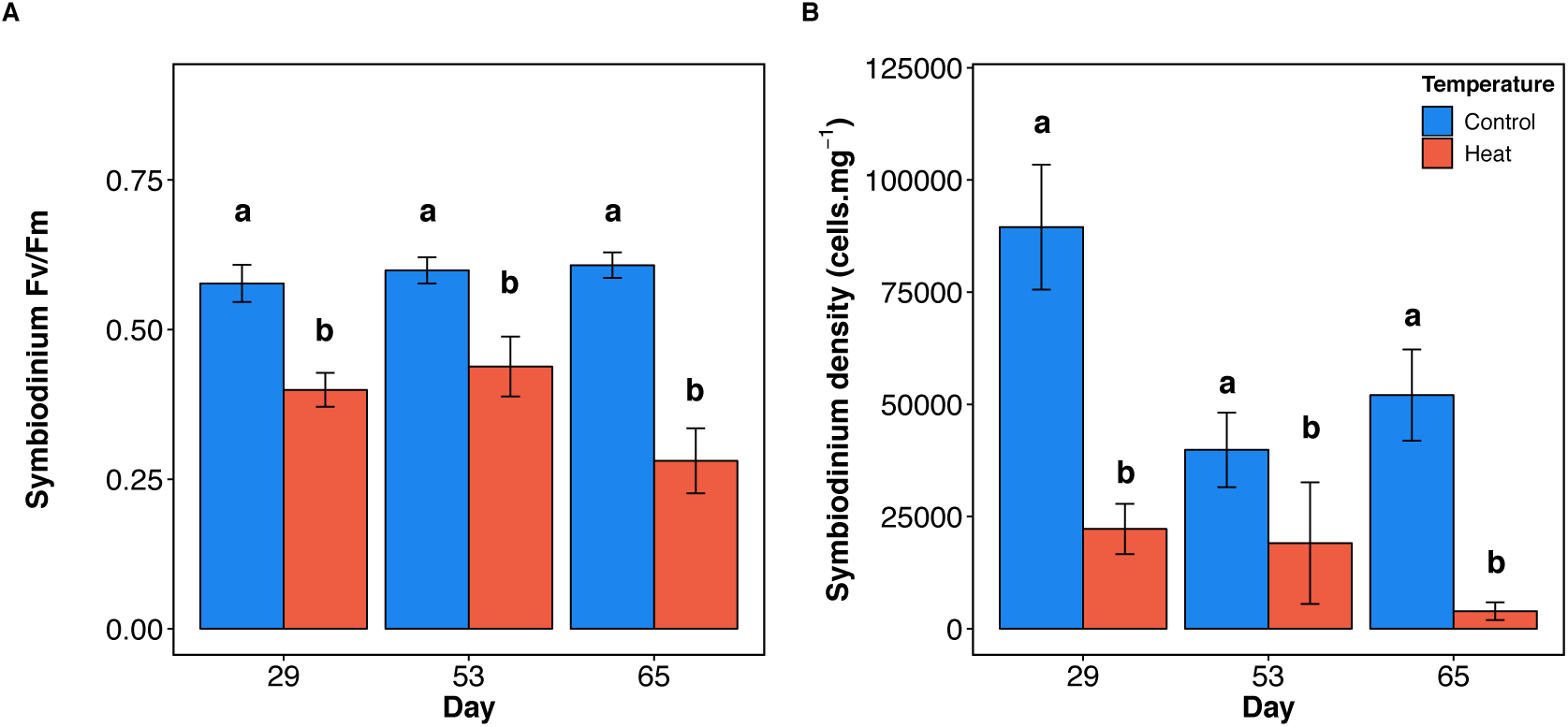
*Symbiodinium* spp. photosynthetic yield and density; Plots represent (A) *Symbiodinium* spp. photosynthetic yield [the maximum dark-adapted yield of photosystem II (Fv/Fm)] and (B) cell density, across 65-day experiment. Lowercase letters represent (Scheirer-Ray-Hare; *p<*0.05). Error bars represent standard error of the mean (n=4).

**Fig. S2:**
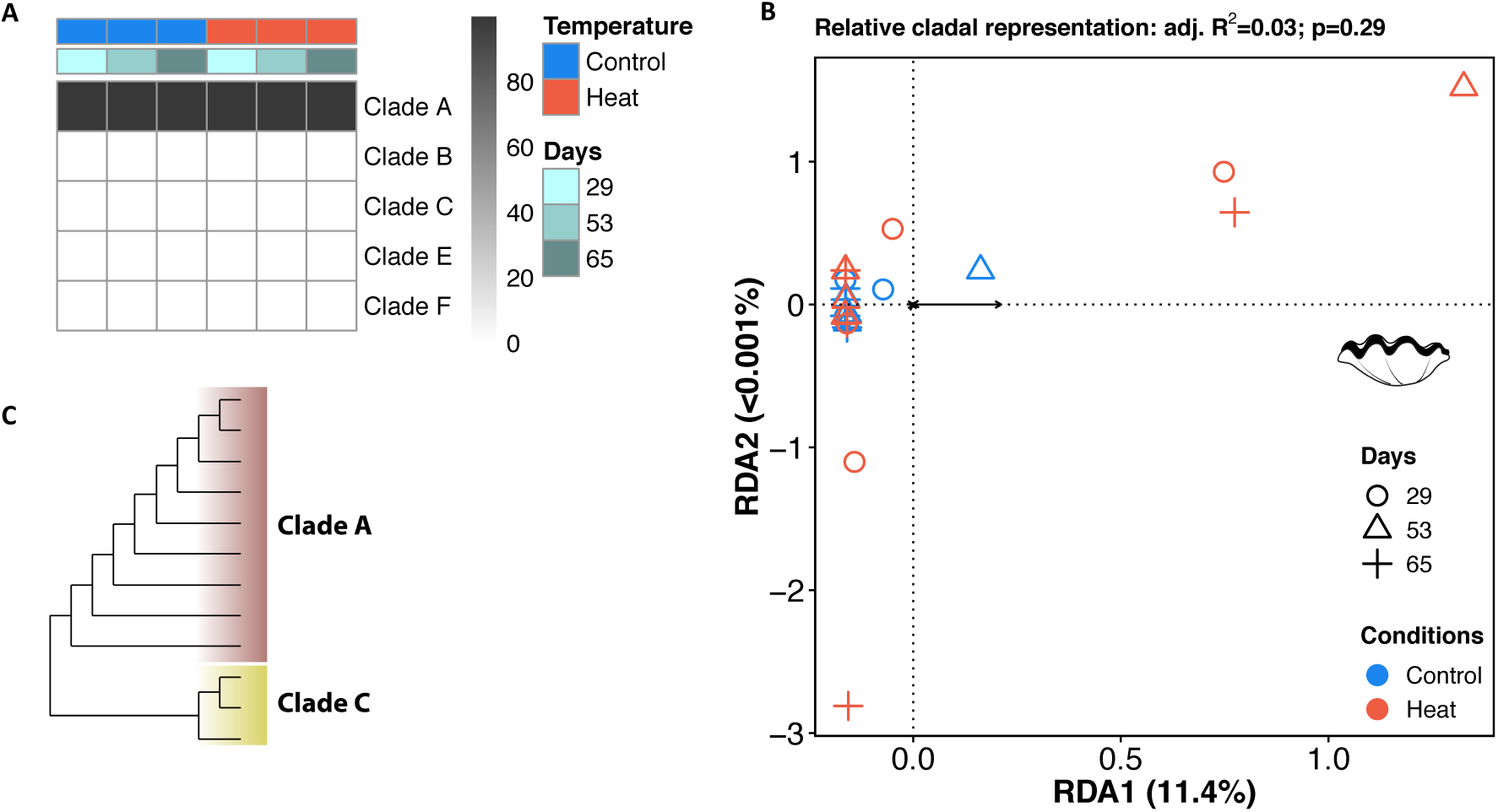
*Symbiodinium* clade representation assessed by qPCR and multivariate analysis. (A) Heatmap showing the median relative clade proportion by group (n=4 individuals/group), as determined by qPCR. (B) RDA representation based on PCoA of Euclidian distances. (C) Cladogram highlighting the diversity of sequences observed for *Symbiodinium* communities in giant clams based on Dada2 clustering and variation of the ITS2 region.

**Fig. S3:**
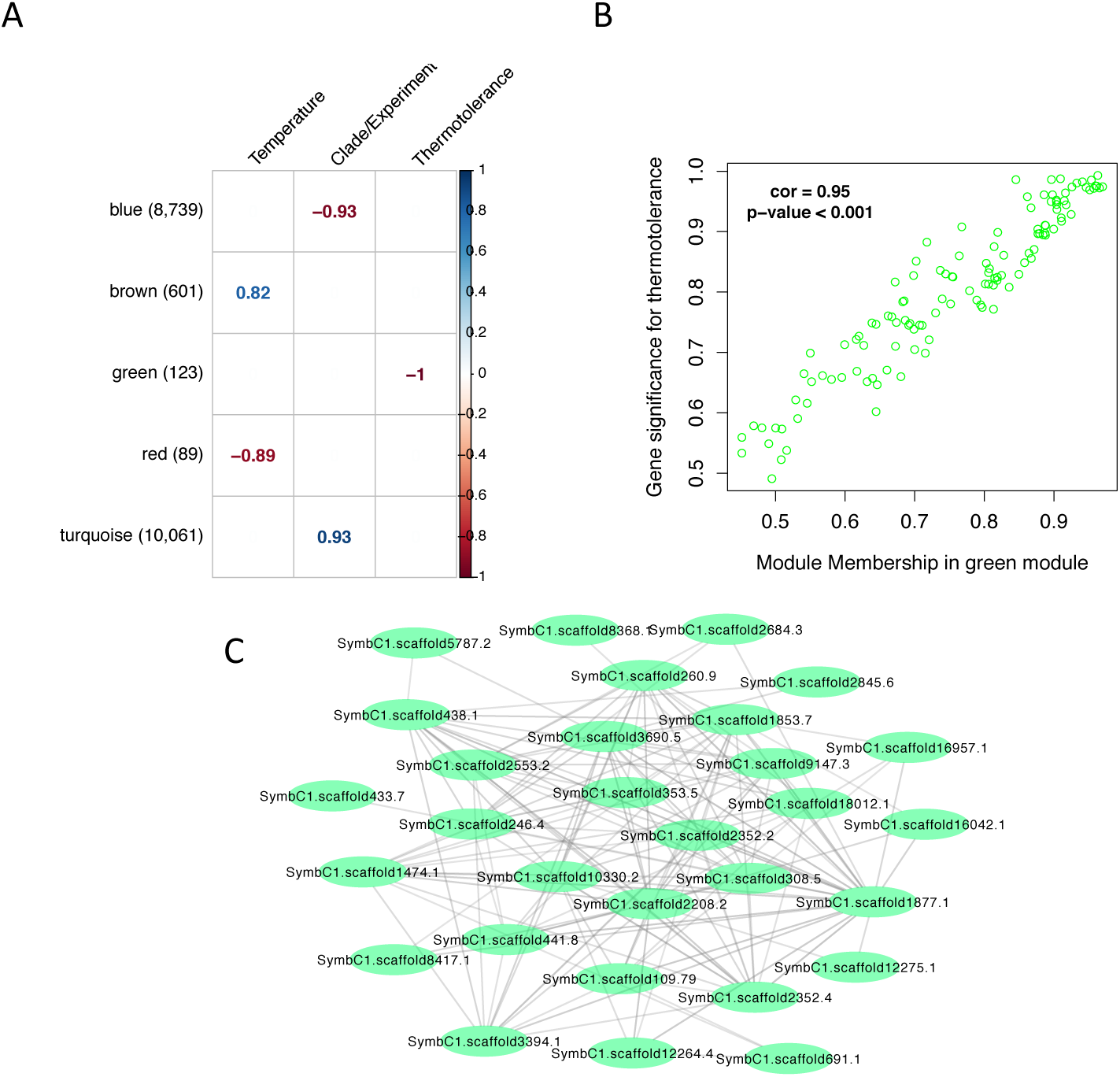
Co-expression analysis of free-living symbionts gene expression based on the *Symbiodinium* clade C1 reference genome (*41*). (A) Correlation matrix of free-living symbionts gene expression against condition factors. Genes are clustered in modules (y–axis) according to their co-expression values. Values in brackets represent the number of genes within each cluster. Values indicate Pearson’s correlation scores and only when significant (*p<*0.01). (B) Plot of gene module membership (kME) against gene significance (GS) value for thermotolerance. Dots represent a single gene. (C) Network representation of the top 30 “hub genes” selected on the kME value for module greenc1

**Table S1: Transcriptome annotations for *Symbiodinium* spp. and *Tridacna maxima*.** Transcripts were aligned against the Uniprot-Swissprot database with BLASTx, and only the best-hit results have been shown.

**Table S2: Excel file featuring gene ontology (GO) enrichment for each module.** Only GO terms with *p<*0.05 and at least three genes were considered to be significantly enriched.

**Table S3: Table S3. Excel file featuring “hub” genes detected within each module.** The top 30 hub genes were selected according to their module membership values (kME).

